# Characterizing Individual and Population-Level Heterogeneity in Infant BMI Trajectories Using Flexible and Functional Modeling Approaches

**DOI:** 10.64898/2026.09.15.751845

**Authors:** Shunzhou Jiang, Alina Yu, Sani M. Roy, Babette B. Zemel, Shana E. McCormack, Rui Xiao

## Abstract

Infant body mass index (BMI) follows a rapid and nonlinear trajectory, yet substantial inter-individual variation complicates accurate characterization of growth dynamics. Here we systematically compared individual- and group-based approaches for modeling longitudinal BMI trajectories in 2,114 healthy infants. Among linear spline, natural cubic spline and fractional polynomial mixed-effects models and SITAR, natural cubic splines provided the best fit while preserving individual variation in trajectory shape. Across all models, males reached their BMI peak earlier than females. To characterize population-level heterogeneity, we further compared latent class mixed models with functional principal component analysis (FPCA) followed by Gaussian mixture clustering. Whereas latent class modeling produced numerous small classes, FPCA identified three interpretable trajectory groups in both sexes, including a rare subgroup characterized by higher BMI and delayed or unestimable peak timing. Maternal gestational diabetes was associated with earlier BMI peak timing in females and was enriched in the rare female trajectory subgroup. Together, these results support combining flexible mixed-effects modeling with functional trajectory analysis to characterize both individual growth dynamics and population-level heterogeneity during infancy.

## Introduction

Early childhood is a critical time period for growth and development, during which physiological systems are rapidly maturing and are sensitive to environmental and prenatal influences^1^. Longitudinal cohort studies with repeated measurements provide an opportunity to determine patterns of change in physiological traits over time. During infancy, these changes are often rapid and highly non-linear, making accurate modeling both crucial and challenging. Body mass index (BMI) across infancy is a clear example. Following birth, BMI increases rapidly, reaching a peak during infancy, and subsequently declines before rising again later in childhood^2^. The shape and timing of these phases vary substantially between individuals, and population-average summaries may conceal differences in growth patterns. Accurately tracking these dynamics thus requires statistical methods that can accommodate non-linearity, irregular measurement timing, and substantial between-individual variability^3^.

Despite the unpredictable nature, growth assessment in clinical settings often relies on overly simplified summaries, such as visual inspection of growth charts. They provide limited insight into individual-level growth patterns, such as rates of change. A range of modeling approaches has thus been applied to childhood BMI data to better characterize growth patterns over a set period of time^4^.

Mixed-effects models are commonly used to analyze repeated BMI measurements, allowing better estimation of a population while accounting for individual-level variation^5^. Spline-based and fractional polynomial formulations enable better modeling of non-linear growth. Non-linear growth models such as SITAR provide an alternative system by modeling differences in growth timing and magnitude across all individuals^6^. Every approach relies on different factors, assumptions, and offers unique advantages and limitations. However, there’s limited information on their performance for modeling infant BMI growth, a period undergoing rapid and heterogeneous change. A main feature of infant BMI trajectories is the timing of the BMI peak, which is the maximum BMI before the decline in early childhood. Estimating this milestone requires well-fitted growth curves^7^; however, understanding how different modeling choices influence the estimation of BMI peak timing and the proportion of children for whom a peak can be reliably identified is an important factor in methodological consideration^8^. In addition to individual-level trajectory modeling, there is importance in identifying subgroups of children who share similar growth patterns^9^. Functional data analysis approaches such as FPCA^10^ offer an alternative method by representing trajectories in a lower-dimensional functional space and clustering individuals based on the prominent variations. Comparing these approaches can help clarify the utility for capturing heterogeneity while still maintaining interpretability, a key advantage of FPCA over LCM.

In this study, we use BMI data from a cohort of healthy infants followed through early childhood to examine the best approaches for modeling infant BMI growth trajectories. The analyses address three main objectives. First, we compare four commonly used individual-level growth modeling approaches (linear spline mixed-effects models, cubic spline mixed-effects models, fractional polynomial mixed-effects models, and SITAR) concerning model fit and the estimation of the timing of the infancy BMI peak. Second, we examine group-based modeling techniques to differentiate subgroups of children with distinct BMI growth patterns. We do this by comparing latent class trajectory models and FPCA-based approaches. Third, we explore the associations between trajectory-derived features such as BMI peak timing and subgroup membership, and demographic factors such as gestational diabetes exposure.

## Methods

### Cohorts and study participants

We analyzed longitudinal infant BMI data collected from subjects participating in “A Study of the Genetic Causes of Complex Pediatric Disorders” at The Children’s Hospital of Philadelphia (CHOP) ^11^. The study was approved by the Institutional Review Board at CHOP, and written informed consent was obtained from participants. Children who had at least two visits in each of the following age ranges (days): 0–88, 89–224, and 225–408 were included in this analysis. Other details of the study and the inclusion and exclusion criteria for this analytic sample were previously published^11^.

The cohort for analysis comprised 2,114 healthy infants, 53% male and 61% African American, with a median follow-up period of 24.93 months. Additional demographic variables, including maternal gestational diabetes and smoking status, were recorded. Infants with less than one year of follow-up were excluded (n=8). Detailed demographic characteristics are presented in **Table 1**, and the observed BMI values and trajectories are shown in **Fig. 1**.

**Table 1.**
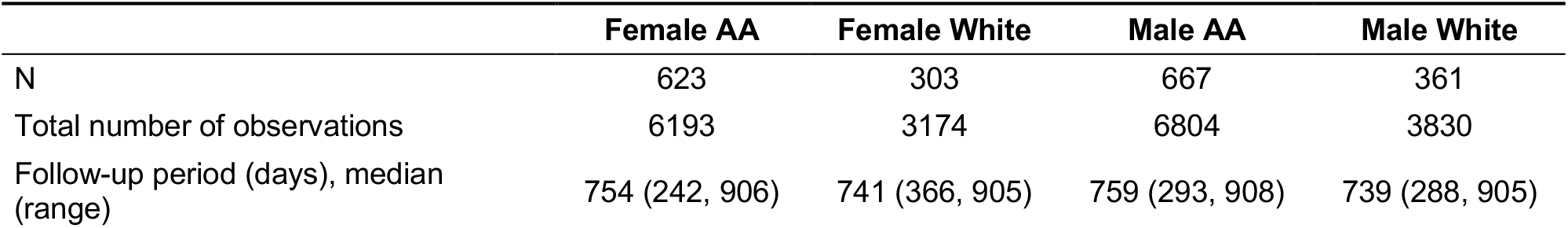
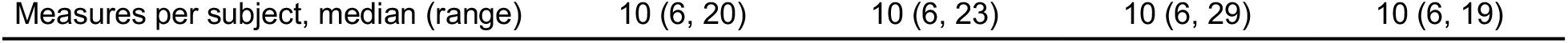
Observed (a) BMI values and (b) BMI trajectories in the cohort, separated by females and males.

**Fig. 1.**
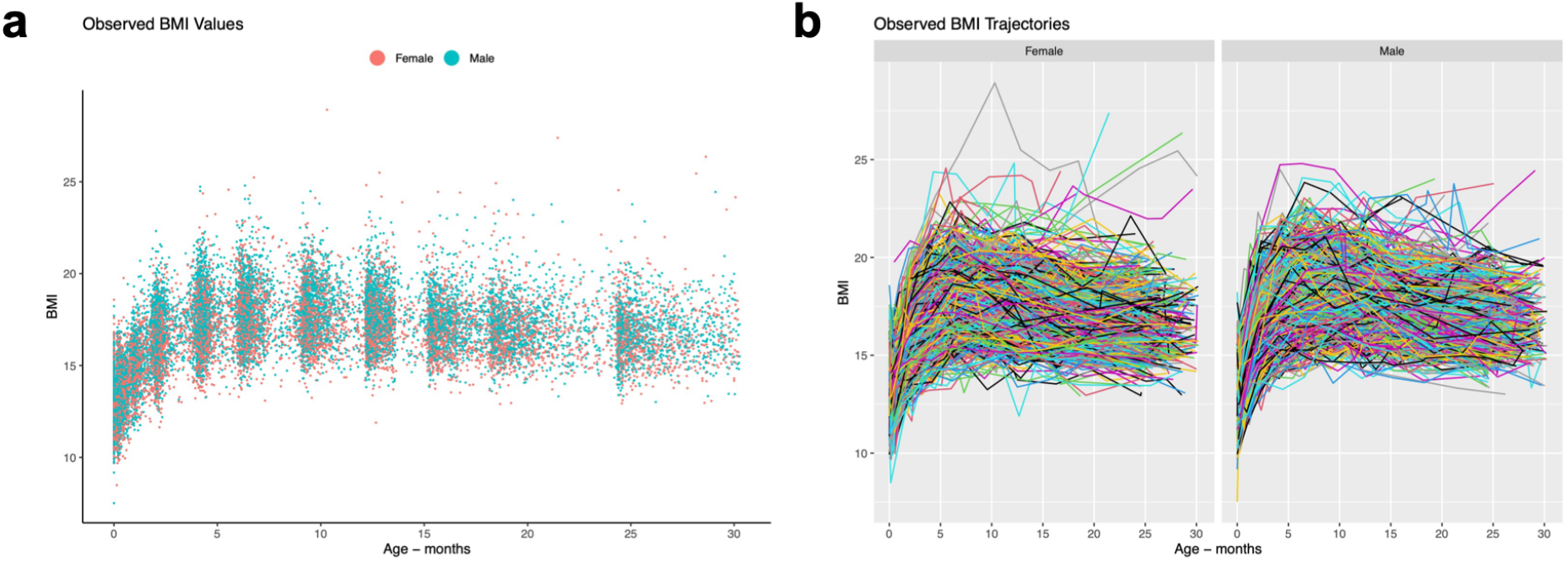
Observed (**a**) BMI values and (**b**) BMI trajectories in the cohort, separated by females and males.

### Individual trajectory modeling

The first aim was to model each infant’s BMI growth trajectory over time. To characterize individual longitudinal patterns, we modeled repeated measurements for each subject using mixed-effects models^12^, which account for both population-level trends (fixed effects) and subject-specific variations (random effects). In general, the mixed-effects model can be written as:

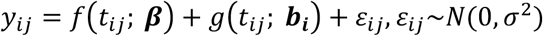

where *y*_*ij*_ denotes the outcome for the individual *i* at time *t*_*ij*_, *f*(*t*_*ij*_; ***β***) represents the fixed effect component describing the overall population trajectory with regression coefficients ***β**, g*(*t*_*ij*_; ***b***_*i*_) captures the random effects of individual *i* with regression coefficients ***b***_*i*_, and *ε*_*ij*_ are residual errors. To flexibly model nonlinear growth patterns, three functional forms of *f*(·) and *g*(·) were employed, including linear spline, natural cubic spline, and fractional polynomial mixed-effects models.

The linear spline model approximates the BMI growth trajectory as a series of connected lines, where *k* knots *κ*_1_, *κ*_2_, …, *κ*_*k*_ are specified and the slope is allowed to change at each knot. In this case, *f*(·) and *g*(·) can be expressed as:

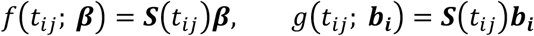

where

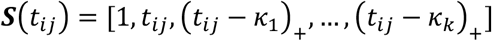

and (*t* − *κ*_*k*_)_+_ = max(0, *t* − *κ*_*k*_). By allowing each segment of the trajectory to own its own slope, linear spline functions capture local changes in growth rate over time. To select the optimal number of knots, candidate values of *k* ranging from 1 to 6 were considered, and the model with the lowest Bayesian Information Criterion (BIC) was selected as the final model.

To capture more complex nonlinear growth patterns, cubic spline mixed-effects models were applied^13^, where the trajectory is represented as a smooth piecewise cubic function of time. Therefore, the fixed- and random-effect components are given by:

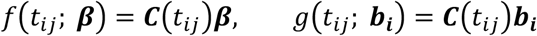

where

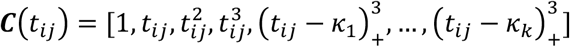

and (*t* − *κ*_*k*_)_+_ = max(0, *t* − *κ*_*k*_). To further avoid high variance of the polynomial functions beyond the boundaries and ensure stable extrapolation, a natural cubic spline was employed, constraining the function to be linear before the first knot and after the last knot. Similar to the linear spline model, candidate values of *k* ranging from 1 to 6 were evaluated, and the optimal model was determined by the lowest BIC value.

To allow more flexible representations of nonlinear time effects, we implemented fractional polynomial mixed-effects models^6^, where the trajectory is expressed as a linear combination of power-transformed terms:

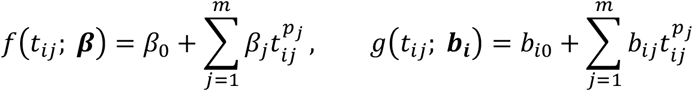

where *m* is the degree of the model, and the powers *p*_*j*_ are selected from a fixed set of 8 candidate values: −2, −1, −0.5, 0.5, 1, 2, and log. A total of 219 candidate models (combinations with *m* ≥ 3) were considered, and the model with the lowest BIC was selected. In our data, the optimal model is:

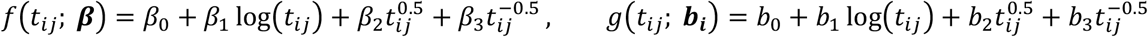

This model was then used for the final trajectory fitting.

Additionally, we considered Super Imposition by Translation and Rotation (SITAR)^14^, a widely used tool for longitudinal trajectory modeling. SITAR represents individual curves as systematically shifted and rescaled versions of a common population mean curve by introducing three subject-specific random effects: size (*α*_*i*_), timing (*β*_*i*_), and velocity (*γ*_*i*_), which account for differences in overall BMI level, growth timing, and growth rate, respectively. It links the outcome *y*_*ij*_ and time *t*_*ij*_ through:

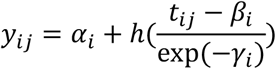

where *h*(·) is a natural cubic spline function representing the fixed-effect population mean trajectory. Through parameter optimization, the goal is to make the individual curves as similar as possible. The optimal number of knots for *h*(·) was determined using the BIC criterion.

We stratified all infants by sex and modeled their growth curves separately using the four methods described above. To compare model performance across these methods, we adopted the Residual Sum of Squares (RSS) between the observed and predicted BMI values as the evaluation metric:

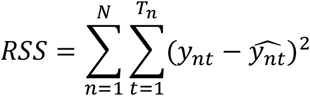

where *N* denotes the total number of infants, *T*_*n*_ is the number of observed time points for subject *n*, while *y*_*nt*_ and 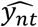 represent the observed and predicted BMI values at time *t* for subject *n*.

### Group-based trajectory modeling

While individual trajectory modeling methods effectively capture both population-level characteristics and individual-level variations, they assume all infants follow a common underlying functional form and differ only through random effects. However, their growth patterns may exhibit systematic heterogeneity and cannot be solely interpreted as random effects. To uncover such latent structures and summarize their high-level growth dynamics, we extended our analysis to group-based trajectory modeling, where infants were clustered into distinct subpopulations that share similar BMI growth profiles.

We first employed the Latent Class Mixed Model (LCMM)^15^, which assumes *G* latent classes and fits a separate mixed-effects model for each class. Conditional on membership in class *g*, the outcome for infant *i* at time *t*_*ij*_ is defined as:

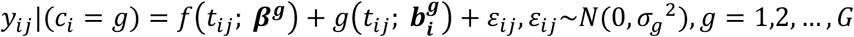

where *f*(*t*_*ij*_; *β*^*g*^) represents the class-specific fixed-effect trajectory, and 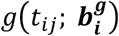 denotes subject-specific random effects within class *g*. After model fitting, each infant is assigned to a latent class *ĝ* with the greatest posterior probability 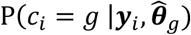, where ***θ***_*g*_ denotes the mixed-effects model parameters for class *ĝ*. The optimal number of latent classes was selected by the BIC evaluated across candidate models with *G* = 2, …,10.

Alternatively, functional principal component analysis (FPCA) can be applied to the full set of infants’ growth trajectories to extract the dominant patterns of variation^10^. Therefore, based on the predicted growth curves from individual trajectory modeling methods, we used FPCA to obtain components that explain 99% of the variation. Each infant receives a score for every principal component, representing the relative contribution of that component to their trajectory. Based on the score matrix, we performed clustering by a Gaussian mixture model (GMM), with the optimal number of clusters determined according to BIC.

Finally, for each method, the subpopulation-level growth trajectory was computed as the average of individual trajectories within each cluster. Group-based trajectory modeling facilitates direct comparisons across subgroups, providing a clearer understanding of heterogeneous growth curve patterns within the cohort.

### BMI peak time identification

After fitting individual trajectory models, BMI was predicted for each infant at equally spaced time points from 0 to 30 months with a 0.1-month interval. The time corresponding to the maximum predicted BMI was regarded as the infant’s peak time. BMI peak times were defined to be estimable if they fell within the 0.5 to 24-month range; otherwise, they were considered as unestimable.

## Results

### Individual trajectory modeling

We stratified infants by their sex and fitted sex-specific BMI trajectory models using linear spline, natural cubic spline, fractional polynomial mixed effects models, and SITAR. Using the fitted models, BMI was predicted for each infant from 0 to 30 months. Population-average BMI growth curves were then computed for females and males, along with their 95% confidence intervals (CIs) (**Figure 2**). Across both sexes and all models, BMI exhibited a unimodal distribution, with an initial rapid increase followed by a gradual decline after the peak. This was further confirmed by the BMI growth velocity of the mean fitted curves (**Figure 3**). Consistent across four methods, over the 30-month period, BMI velocity was highest immediately after birth, steadily decreased over time, became negative, and eventually approached zero. We next identified the BMI peak time for each infant and compared the peak time distributions between females and males (**Figure 4, Table 2**). Across all methods, males exhibited significantly earlier BMI peaks than females, as determined by two-sided t-tests.

**Table 2.** Observed (a) BMI values and (b) BMI trajectories in the cohort, separated by females and males.

|  | Female | Male |
| --- | --- | --- |
| Method 1: Linear spline mixed-effects model (knots selected by BIC) | 3908.7 | 5234.7 |
| Method 2: Cubic spline mixed-effects model (knots selected by BIC) | 2926.5 | 3389.5 |
| Method 3: Fractional polynomial mixed-effects model, $\ln(t) + t^{-1/2} + t^{1/2}$ | 4455.2 | 5631.8 |
| Method 4: SITAR (log-transformed month; df selected by BIC) | 4504.3 | 5674.8 |

**Fig. 2.**
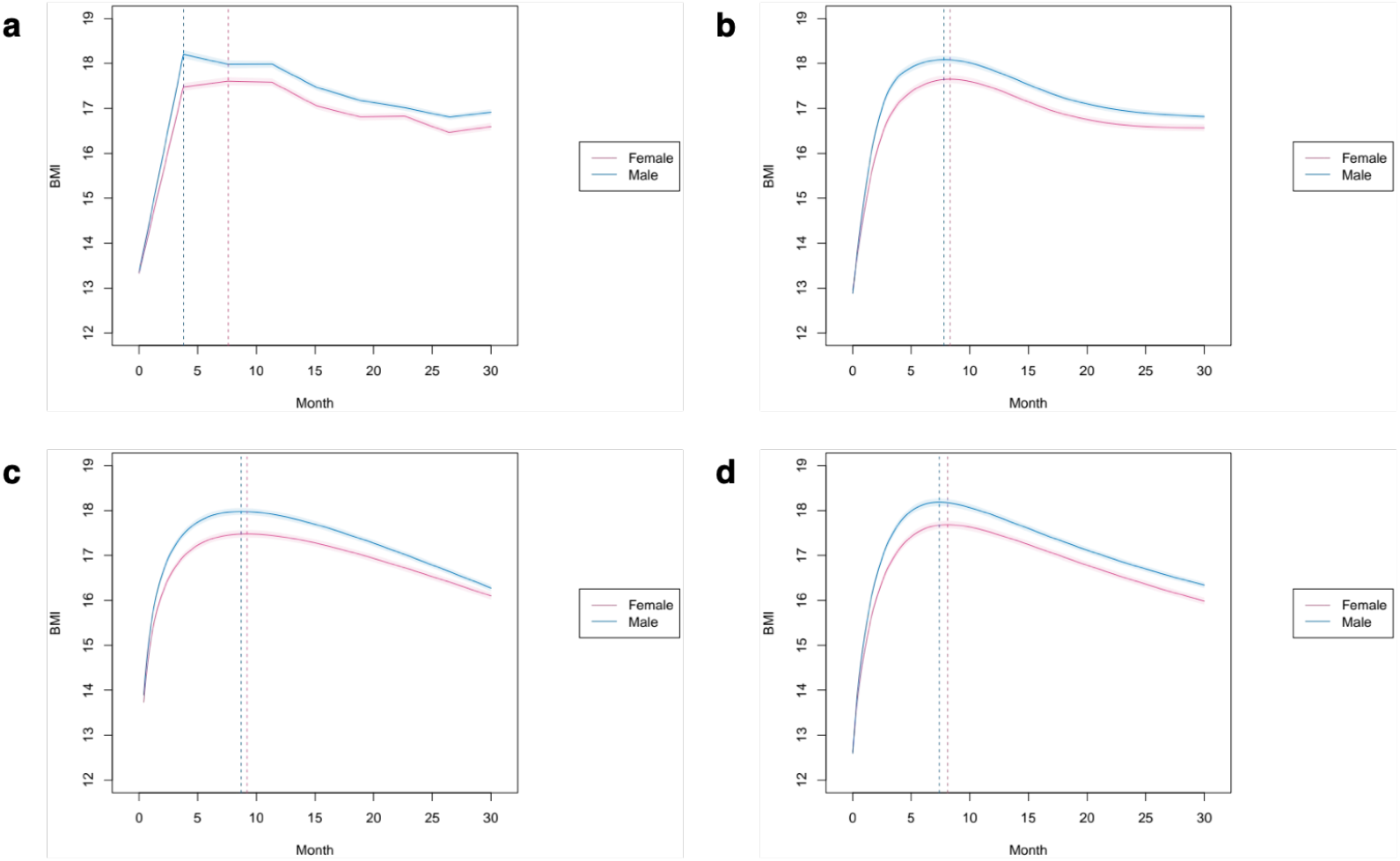
Population-average trajectories with 95% CIs for females (in purple) and males (in blue) from (**a**) linear spline mixed-effects model, (**b**) natural cubic spline mixed-effects model, (**c**) fractional polynomial mixed-effects model, (**d**) SITAR. Dashed lines indicate the BMI peak times of the population-average trajectories.

**Fig. 3.**
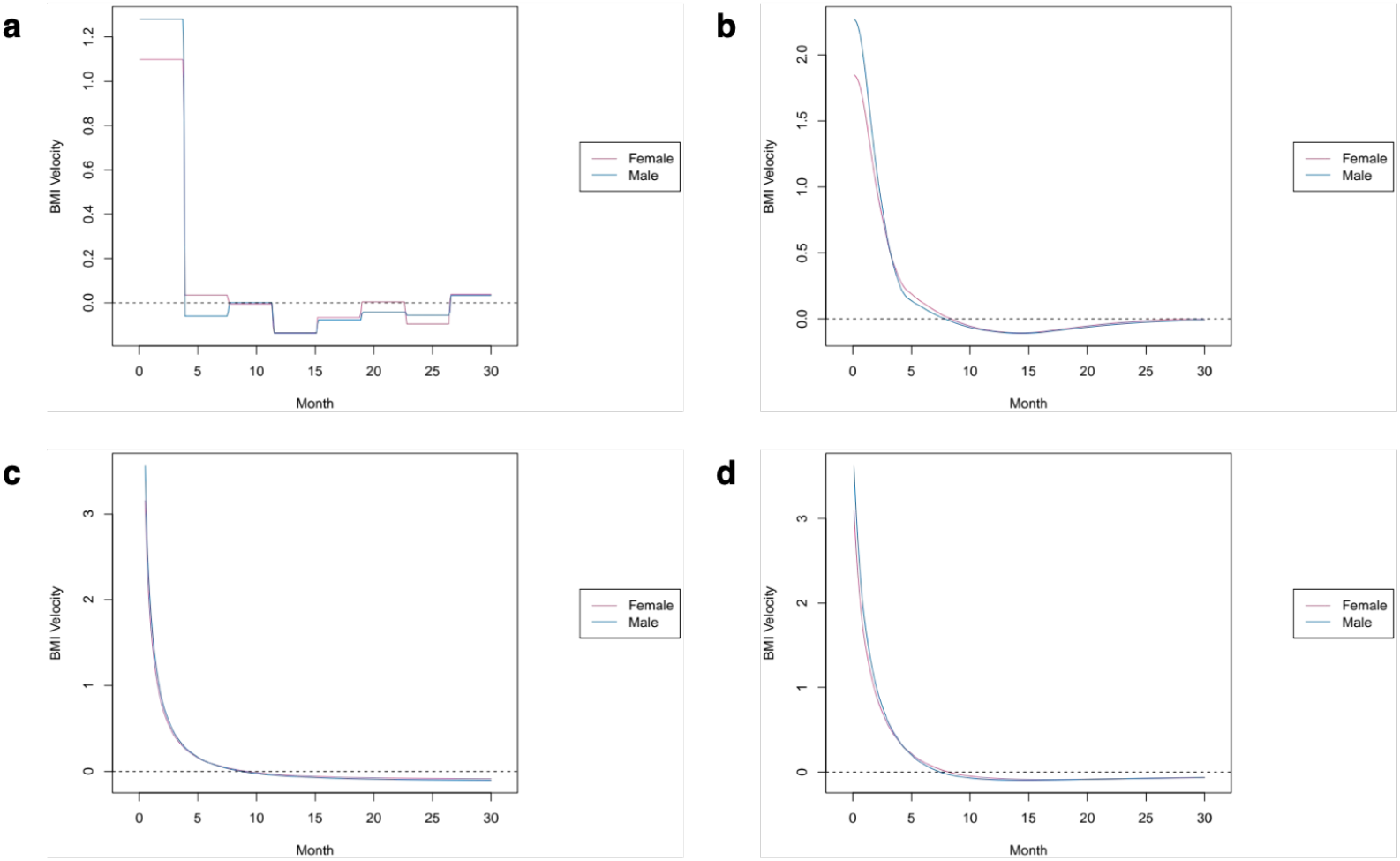
BMI growth velocity of the population-average trajectories for females (in purple) and males (in blue) from (**a**) linear spline mixed-effects model, (**b**) natural cubic spline mixed-effects model, (**c**) fractional polynomial mixed-effects model, (**d**) SITAR. Dashed lines indicate zero growth velocity.

**Fig. 4.**
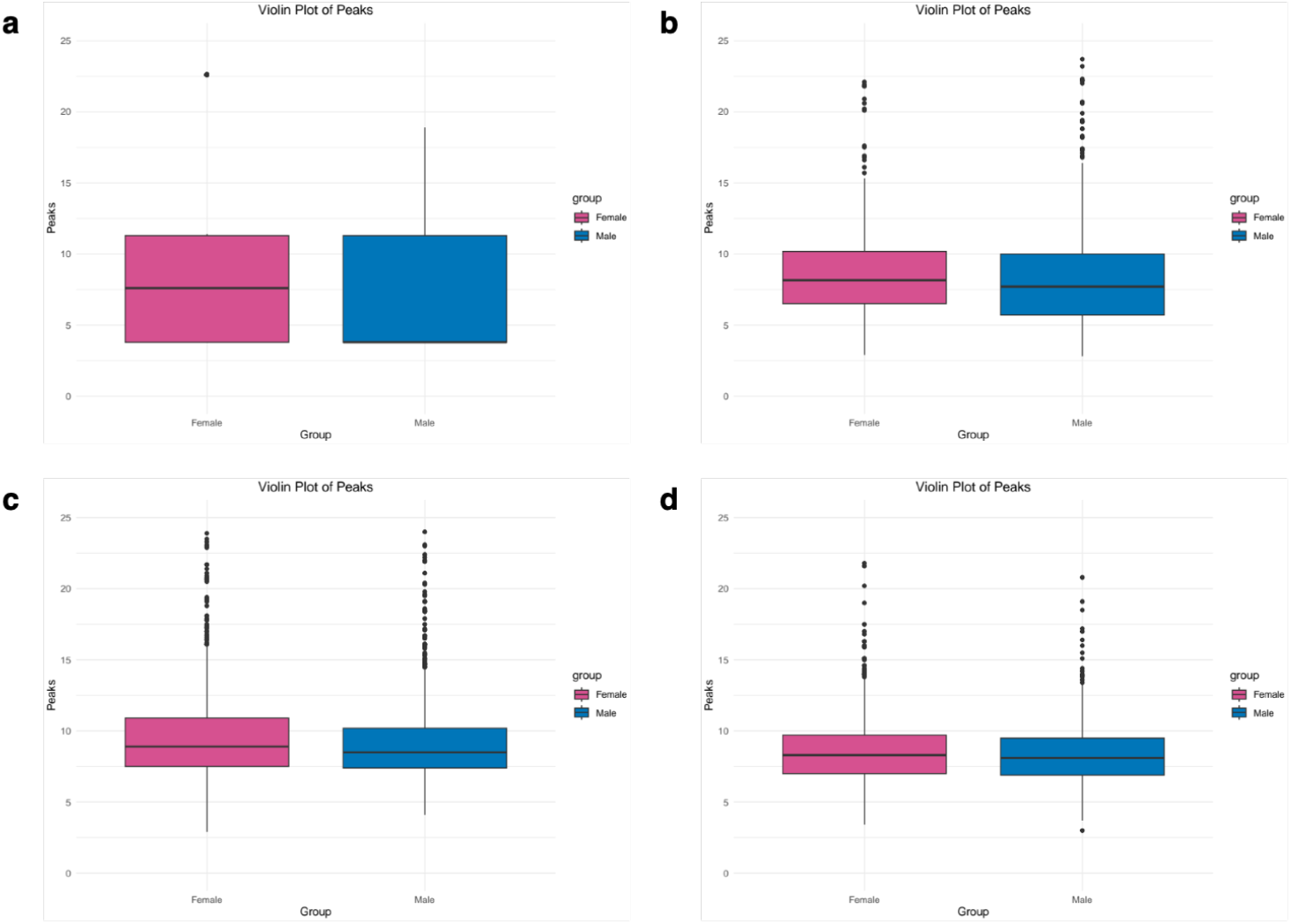
Boxplots displaying the individual peak time distribution for females (in purple) and males (in blue) from (**a**) linear spline mixed-effects model, (**b**) natural cubic spline mixed-effects model, (**c**) fractional polynomial mixed-effects model, (**d**) SITAR.

To benchmark the predictive performance of these four models, we calculated the RSS between observed and predicted BMI values. In both sexes, the natural cubic spline mixed-effect model achieved the lowest RSS, demonstrating its superior predictive performance (**Table 3**). Visualization of individual predicted BMI curves (**Supplementary Fig. 1**) further confirmed that this model preserved inter-individual variability and best captured observed BMI patterns. In contrast, the linear cubic spline mixed-effect model imposed rigid piecewise linearity, while the fractional polynomial mixed-effects model and SITAR tended to produce overly smooth and homogeneous curves across individuals, lacking variability. Collectively, the natural cubic spline mixed-effect model yielded the best performance both quantitatively and visually. Thus, predicted growth curves from this model were used as input for subsequent group-based trajectory analyses.

**Table 3.** Individual peak time statistics (mean, standard deviation, and fraction of estimable peaks) for females and males based on the observed data, linear spline mixed-effects model, natural cubic spline mixed-effects model, fractional polynomial mixed-effects model, and SITAR. P-values of two-sided t-test comparing mean peak times between females and males for each method are also reported.

|  | Observed data | Method 1: Linear spline LME | Method 2: Cubic spline LME | Method 3: Fractional polynomial LME | Method 4: SITAR |
| --- | --- | --- | --- | --- | --- |
| Female | 9.49 (4.40) (93.98%) | 7.76 (4.13) (94.08%) | 8.35 (2.81) (93.58%) | 9.64 (3.13) (95.29%) | 8.59 (2.27) (99.80%) |
| Male | 8.99 (4.41) (96.21%) | 6.58 (3.62) (95.49%) | 7.99 (3.37) (97.48%) | 9.17 (2.72) (97.75%) | 8.30 (2.05) (100%) |
| P-value | 0.005 | $1.824 \times 10^{-11}$ | 0.008 | 0.0003 | 0.0002 |

**Supplementary Fig. 1.**
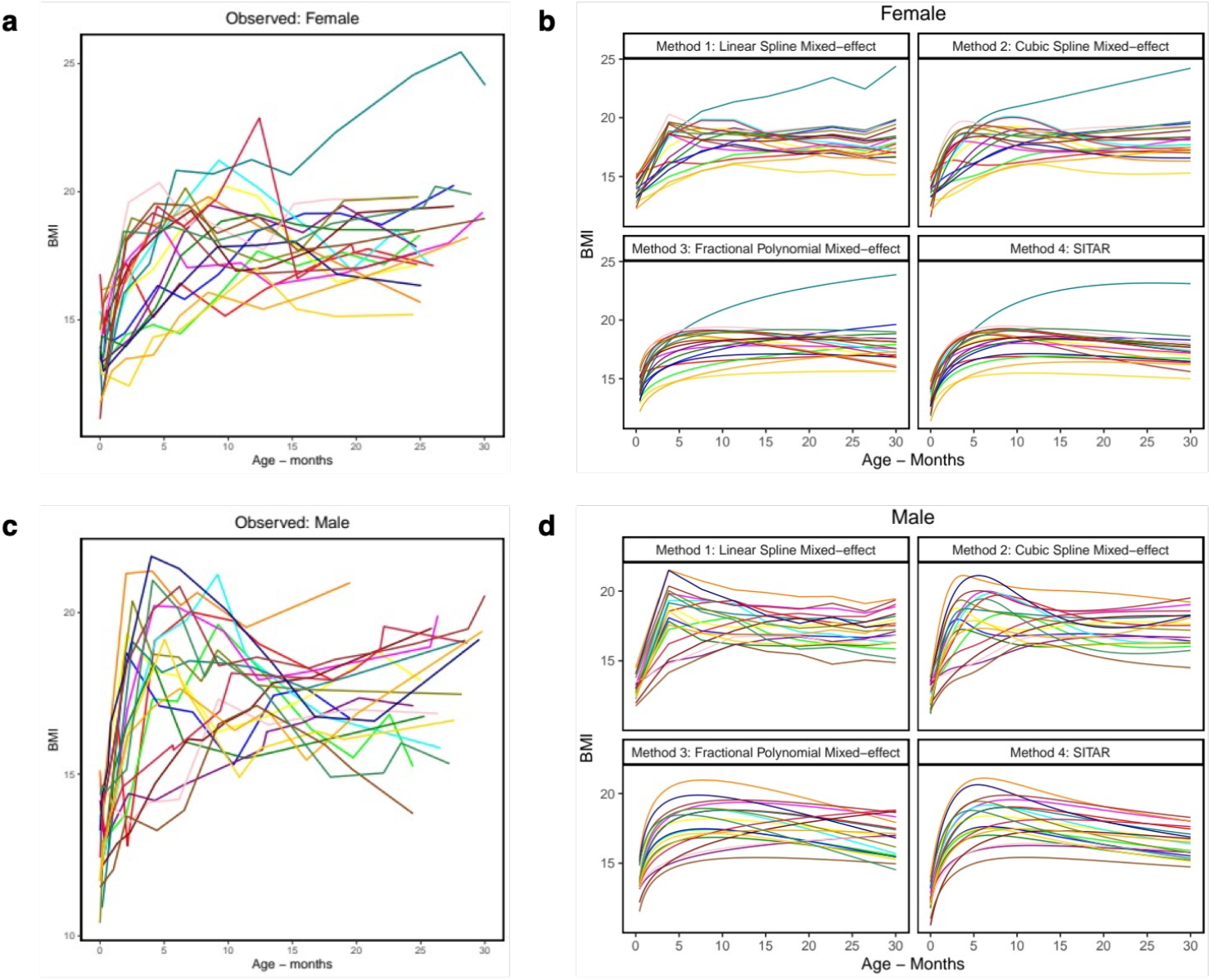
Individual fitted BMI curves for 20 randomly selected children per sex.

### Group-based trajectory modeling

To model the intrinsic heterogeneity in infant growth dynamics, we first applied LCMM to females and males, separately. Based on the latent class memberships estimated by LCMM and individual BMI trajectories predicted from the natural cubic spline model, we visualized class-specific mean growth curves along with their 95% CIs (**Figure 5**). According to the BIC criterion, LCMM identified 10 latent classes for females and 9 for males, some of which contained only a very small fraction of infants. While capturing fine-grained variations, many identified patterns were overly specific, lacked generalizability, and were difficult to interpret. To derive more robust and interpretable subpopulation structures, we performed FPCA on the predicted BMI trajectories, followed by GMM clustering on the FPCA scores. BIC identified three optimal classes for both females and males, each cluster representing a reasonable fraction of the cohort and reflecting distinct yet generalizable growth patterns (**Figure 6**). Notably, a rare class was observed in both sexes (Class 2 for females and Class 1 for males), characterized by higher BMI and delayed BMI peak times. We further compared the individual peak time distribution across the three clusters for each sex. Infants belonging to this rare class had a larger proportion of unestimable peaks (i.e., peak time > 24 months) and significantly delayed peak times among those with estimable peaks (**Figure 7**), as confirmed by the statistical test of equal proportions and two-sided t-tests across the three classes.

**Fig. 5.**
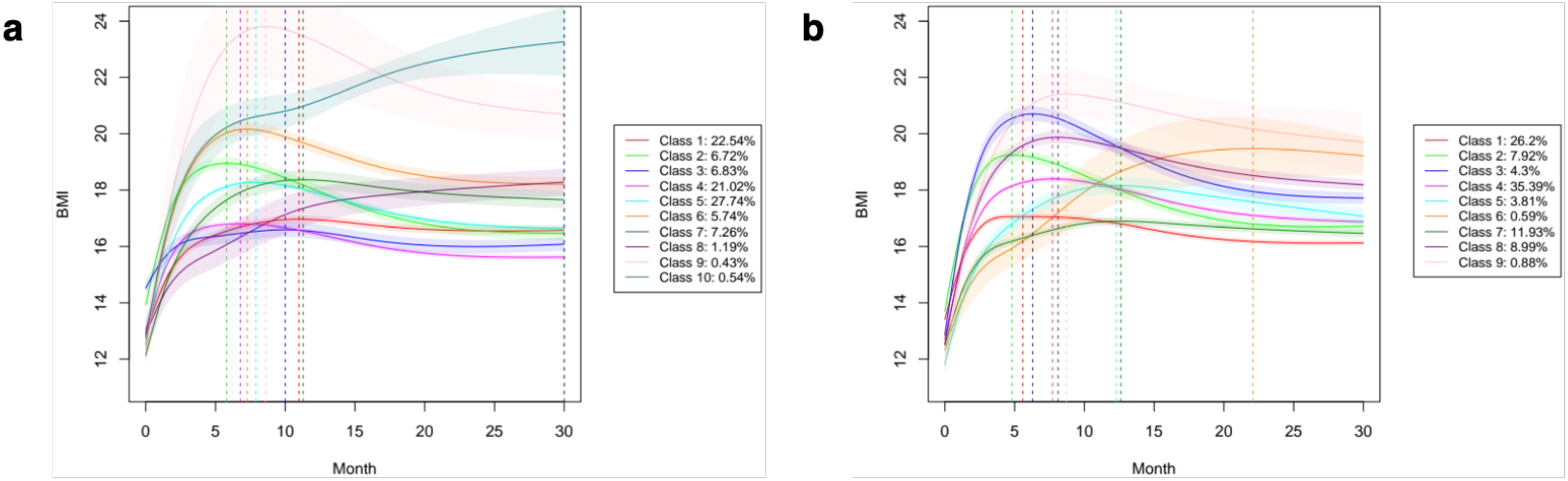
Class-specific mean growth curves with 95% CIs for LCMM-derived clusters, where mean curves were obtained by averaging individual trajectories predicted from the natural cubic spline mixed-effects model, shown for (**a**) females and (**b**) males. The proportion of infants in each cluster is indicated in the figure legend.

**Fig. 6.**
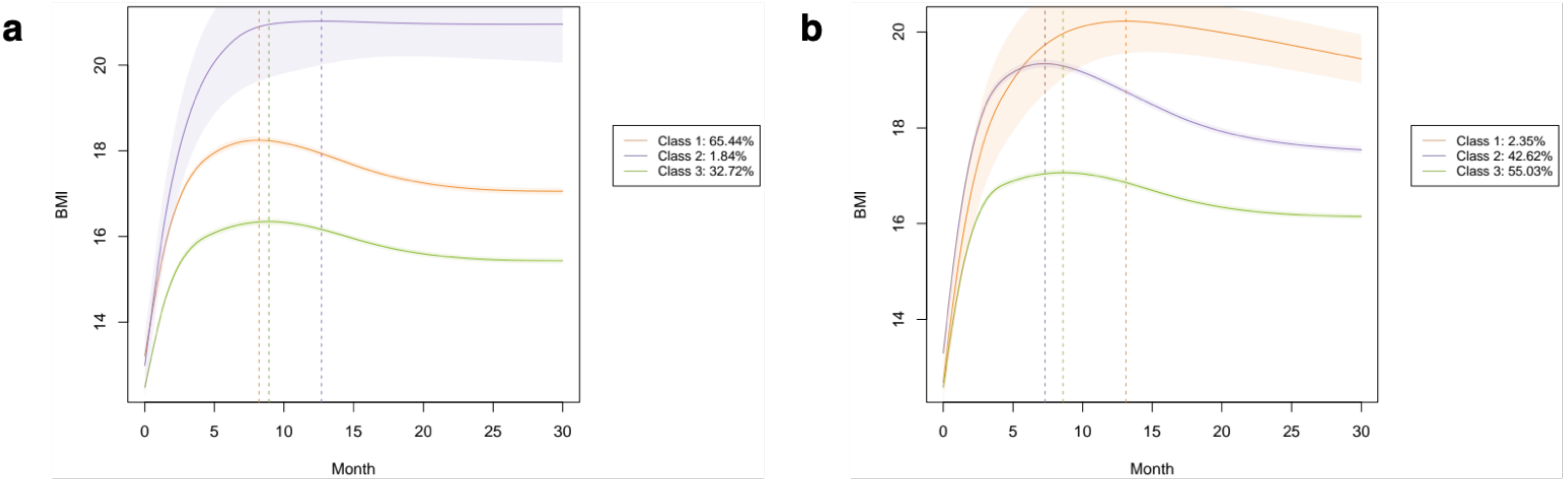
Class-specific mean growth curves with 95% CIs for FPCA-derived clusters, where mean curves were obtained by averaging individual trajectories predicted from the natural cubic spline mixed-effects model, shown for (**a**) females and (**b**) males. The proportion of infants in each cluster is indicated in the figure legend.

**Fig. 7.**
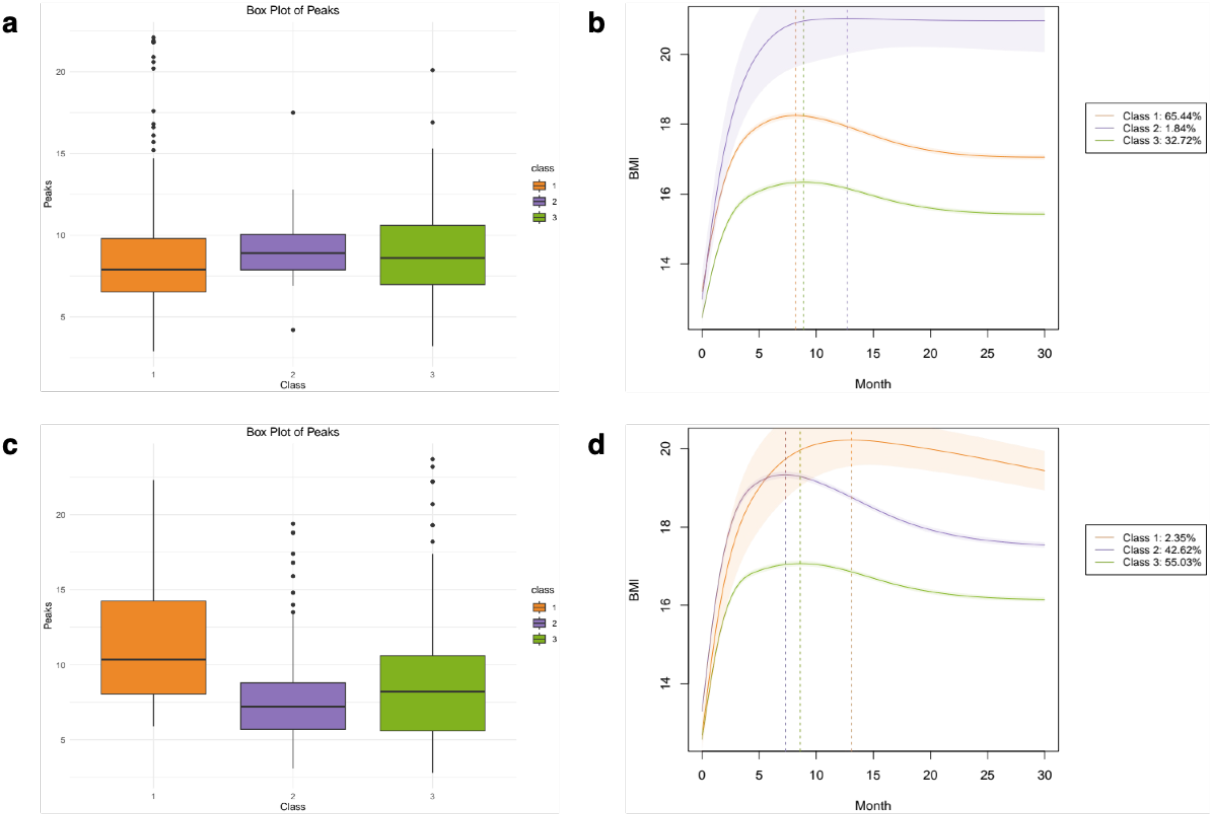
Boxplots displaying the individual peak time distributions across FPCA-derived clusters for (**a**) females and (**b**) males.

### Association analysis with demographic variables

Having characterized individual-level and group-level BMI growth trajectories, we next investigated how these characteristics were associated with phenotypic variables, including maternal gestational diabetes and smoking status. First, we visualized the individual peak time distribution (**Figure 8**) and tested whether the mean estimable peak times differed across levels of demographic variables. As shown in **Table 4**, female infants born to mothers with gestational diabetes reached their BMI peak significantly earlier, whereas no significant difference was observed among male infants. Next, we linked the FPCA-identified clusters with the demographic characteristics by quantifying the fraction of infants exposed to gestational diabetes or maternal smoking within each cluster. **Table 5** demonstrates that in females, infants in Class 2, the rare subgroup with higher BMI and delayed peak times, were more likely to be exposed to gestational diabetes, while no significant association was detected for the smoking status.

**Table 4.** Fraction of estimable peaks and mean peak times stratified by gestational diabetes and smoking status for females and males. P-values of two-sided t-tests comparing mean peak times between levels of each demographic variable are also reported.

|  | Gestational diabetes |  |  | Smoker in house |  |  |
| --- | --- | --- | --- | --- | --- | --- |
|  | Y | N | P-value | Y | N | P-value |
| Female | 7.21<br>(77.42%) | 8.45<br>(93.94%) | 0.005 | 8.47<br>(95.40%) | 8.40<br>(92.80%) | 0.77 |
| Male | 7.33<br>(95.83%) | 8.14<br>(97.60%) | 0.24 | 8.14<br>(98.10%) | 8.11<br>(97.40%) | 0.93 |

**Table 5.**
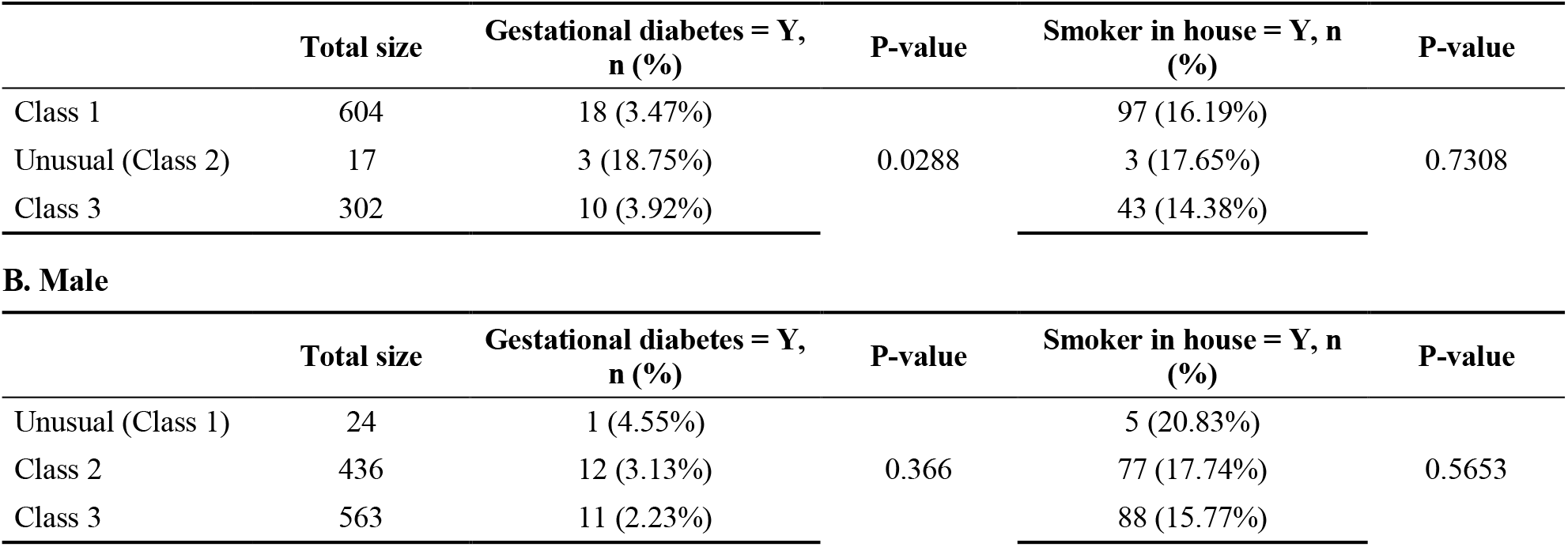
Fraction of infants exposed to gestational diabetes and maternal smoking in each FPCA-derived cluster. P-values testing the equality of proportions are also reported.

A. Female
|  | Total size | Gestational diabetes = Y, n (%) | P-value | Smoker in house = Y, n (%) | P-value |
| --- | --- | --- | --- | --- | --- |
| Class 1 | 604 | 18 (3.47%) | 0.0288 | 97 (16.19%) | 0.7308 |
| Unusual (Class 2) | 17 | 3 (18.75%) |  | 3 (17.65%) |  |
| Class 3 | 302 | 10 (3.92%) |  | 43 (14.38%) |  |

|  | Total size | Gestational diabetes = Y, n (%) | P-value | Smoker in house = Y, n (%) | P-value |
| --- | --- | --- | --- | --- | --- |
| Unusual (Class 1) | 24 | 1 (4.55%) | 0.366 | 5 (20.83%) | 0.5653 |
| Class 2 | 436 | 12 (3.13%) |  | 77 (17.74%) |  |
| Class 3 | 563 | 11 (2.23%) |  | 88 (15.77%) |  |

**Fig. 8.**
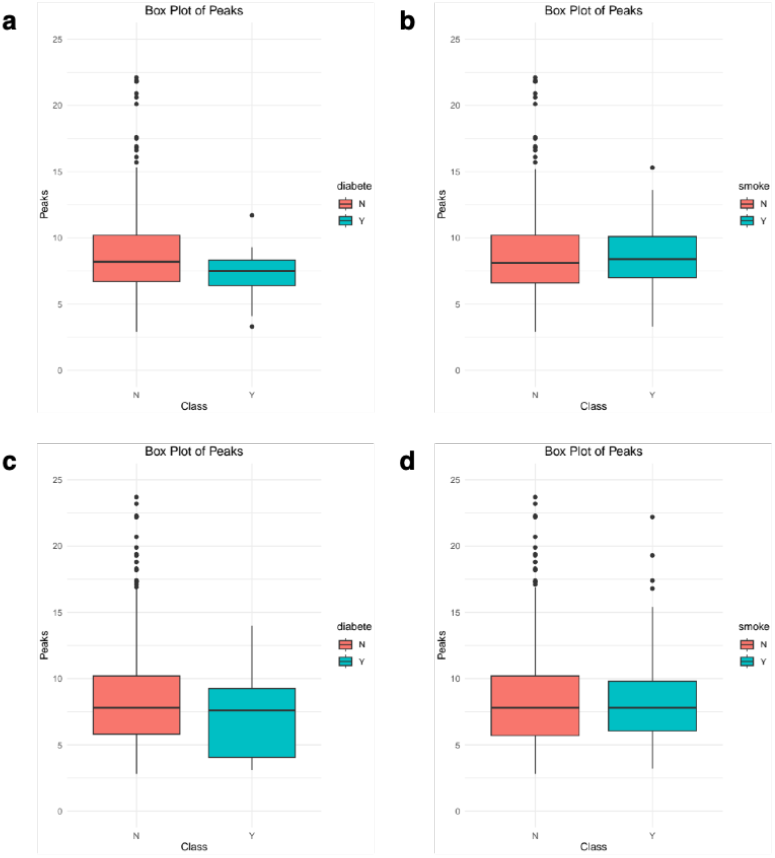
Boxplots displaying the individual peak time distributions separated by demographic variables including gestational diabetes and smoking status for (**a**) females and (**b**) males.

## Discussion

In this study, we employed four individual and two group-based trajectory modeling methods to model the growth characteristics of infant BMI. Individual trajectory modeling methods included linear spline mixed-effects model, natural cubic spline mixed-effects model, fractional polynomial mixed-effects model, and SITAR. Through qualitative and quantitative evaluations, the natural cubic spline mixed-effects model outperformed others by achieving the best prediction accuracy and preserving inter-individual variability in growth trajectories. Across all methods, male infants consistently peaked earlier than females across all methods. Additionally, to capture population-level heterogeneity, we applied LCMM and FPCA to identify the latent cluster membership and group infants with similar BMI growth characteristics. While LCMM tended to overcluster the data, FPCA generated more biologically meaningful results and clustered both females and males into three clusters, with peak time distributions significantly different among them. Interestingly, in both sexes, one rare cluster was characterized by elevated BMI levels and delayed peak timing during infancy. Further association analyses between demographic variables revealed that female infants exposed to gestational diabetes had significantly earlier peaks, and those belonging to the high-BMI, delayed-peak cluster (Class 2) were more likely to have been exposed to gestational diabetes.

Several limitations should be considered. The analysis was conducted in a single cohort, and both the preferred modeling strategy and the identified trajectory subgroups should be evaluated in independent populations. Measurement schedules and follow-up duration also varied across infants, which may increase uncertainty in trajectory estimates near the boundaries of the observation period. In particular, classification of peaks after 24 months as unestimable limits interpretation of very late-peaking trajectories. In addition, the trajectory clusters should be viewed as useful summaries of growth heterogeneity rather than evidence for intrinsically discrete biological populations, as some variation in infant growth is likely continuous. The relatively small size of the rare trajectory groups further limits inference regarding their demographic or clinical correlates.

Overall, our results show that methodological choices can substantially influence the characterization of infant BMI growth. Flexible mixed-effects modeling provides accurate reconstruction of nonlinear individual trajectories, whereas FPCA-based analysis offers an interpretable representation of population-level heterogeneity. Combining these two levels of analysis enables both robust estimation of dynamic features such as BMI peak timing and identification of distinct longitudinal growth patterns. Future studies in larger and independent cohorts could determine whether these trajectory-derived phenotypes are associated with later childhood adiposity, metabolic outcomes or other developmental characteristics.

## References

1 Dietz, W. H. Critical periods in childhood for the development of obesity. Am J Clin Nutr 59, 955–959 (1994). 10.1093/ajcn/59.5.955

2 Rolland-Cachera, M. F. et al. Adiposity rebound in children: a simple indicator for predicting obesity. Am J Clin Nutr 39, 129–135 (1984). 10.1093/ajcn/39.1.129

3 Tu, Y. K., Tilling, K., Sterne, J. A. & Gilthorpe, M. S. A critical evaluation of statistical approaches to examining the role of growth trajectories in the developmental origins of health and disease. Int J Epidemiol 42, 1327–1339 (2013). 10.1093/ije/dyt157

4 Tilling, K., Macdonald-Wallis, C., Lawlor, D. A., Hughes, R. A. & Howe, L. D. Modelling childhood growth using fractional polynomials and linear splines. Ann Nutr Metab 65, 129–138 (2014). 10.1159/000362695

5 Laird, N. M. & Ware, J. H. Random-effects models for longitudinal data. Biometrics 38, 963–974 (1982).

6 Wen, X., Kleinman, K., Gillman, M. W., Rifas-Shiman, S. L. & Taveras, E. M. Childhood body mass index trajectories: modeling, characterizing, pairwise correlations and socio-demographic predictors of trajectory characteristics. BMC Med Res Methodol 12, 38 (2012). 10.1186/1471-2288-12-38

7 Silverwood, R. J., De Stavola, B. L., Cole, T. J. & Leon, D. A. BMI peak in infancy as a predictor for later BMI in the Uppsala Family Study. Int J Obes (Lond) 33, 929–937 (2009). 10.1038/ijo.2009.108

8 Bornhorst, C. et al. Potential selection effects when estimating associations between the infancy peak or adiposity rebound and later body mass index in children. Int J Obes (Lond) 41, 518–526 (2017). 10.1038/ijo.2016.218

9 Nagin, D. S. & Odgers, C. L. Group-based trajectory modeling in clinical research. Annu Rev Clin Psychol 6, 109–138 (2010). 10.1146/annurev.clinpsy.121208.131413

10 Chen, A., Stein, R., Baldassano, R. N. & Huang, J. Learning Longitudinal Patterns and Subtypes of Pediatric Crohn Disease Treated With Infliximab via Trajectory Cluster Analysis. J Pediatr Gastroenterol Nutr 74, 383–388 (2022). 10.1097/MPG.0000000000003370

11 Roy, S. M. et al. Body mass index (BMI) trajectories in infancy differ by population ancestry and may presage disparities in early childhood obesity. J Clin Endocrinol Metab 100, 1551–1560 (2015). 10.1210/jc.2014-4028

12 Johnson, W., Balakrishna, N. & Griffiths, P. L. Modeling physical growth using mixed effects models. Am J Phys Anthropol 150, 58–67 (2013). 10.1002/ajpa.22128

13 Burrows, K. et al. A framework for conducting GWAS using repeated measures data with an application to childhood BMI. Nat Commun 15, 10067 (2024). 10.1038/s41467-024-53687-3

14 Cole, T. J., Donaldson, M. D. & Ben-Shlomo, Y. SITAR--a useful instrument for growth curve analysis. Int J Epidemiol 39, 1558–1566 (2010). 10.1093/ije/dyq115

15 Proust-Lima, C., Philipps, V. & Liquet, B. Estimation of Extended Mixed Models Using Latent Classes and Latent Processes: The R Package lcmm. Journal of Statistical Software 78, 1–56 (2017). 10.18637/jss.v078.i02

